# The Response of Culturally Important Plants to Experimental Warming and Grazing in Pakistan Himalayas

**DOI:** 10.1101/2020.08.07.241117

**Authors:** Saira Karimi, Muhammad Ali Nawaz, Saadia Naseem, Ahmed Akrem, Olivier Dangles, Zahid Ali

**Affiliations:** Department of Biosciences, COMSATS University Islamabad (CU), Islamabad, Pakistan; Department of Animal Sciences, Quaid-i-Azam University, Islamabad, Pakistan; Department of Botany, Institute of Pure and Applied Biology, Bahauddin Zakariya University, Multan, Pakistan; CEFE, UMR 5175, CNRS, Université de Montpellier, Université Paul Valéry Montpellier, EPHE, IRD, Montpellier, France

**Keywords:** Climate change, indigenous people, Khunjerab National Park, medicinal plants, open-top chambers

## Abstract

The response of wild plants towards climate warming is taxa specific, but overgrazing could also be a determining factor for the alpine ecosystems. Overgrazing and climate warming are important drivers of alpine grassland degradation worldwide. Local indigenous peoples will be the first impacted by such degradation due to impacts on animal production and the availability of local medicinal plants. Studies on plant responses to overgrazing and climate change are rarely performed to assess threats to these biological and cultural systems. Long-term observations or manipulative experiments are promising, but rarely use strategies to evaluate the sensitivity and vulnerability of such ecosystems to climatic change. We studied the combined effects of overgrazing and increased temperatures on culturally important medicinal plants of Khunjerab National Park, Pakistan. Three experimental treatments were used (control, warming through an open-top chamber, and exclusion of grazing animals vs. the control). These experimental plots were installed at different elevations (3352-4969 m) and were monitored routinely. Grazing reduced vegetation cover & biomass by 2.3% and 6.26%, respectively, but that was not significant due to the high variability among study plots. However, warming significantly increased the overall percentage cover and biomass of all target plant species, ranging from 1±0.6% in *Bistorta officinalis* to 18.7 ± 4.9% in *Poa alpina*. Thus, warming may increase the availability of therapeutic plants for indigenous people while overgrazing would have deteriorating effects locally. This research illustrates that vegetation sensitivity to warming and overgrazing is likely to affect man– environment relationships, and traditional knowledge on a regional scale.

## Introduction

Climate change is altering the structure and function of high elevation ecosystems. Mountains are amazingly diverse ecosystems that hold high proportions of endemic species [1–5]. These systems are particularly vulnerable to climate change with most species distribution models projecting drastic changes in community composition and distribution range[6]. Importantly, climate change impacts high altitude regions by a combination of stressors such as land and water degradation due to farming [7]. These stressors have profound effects on the habitability of the mountainous ecosystem and the human communities that depend on them (IPCC 2019), in the inter-tropical region (Himalayas, Andes, Eastern Africa). In the Himalayas, an increase of 3.7 °C is expected in global mean surface temperatures between 2018–2100 (relative to 1986–2005). That is expected to cause substantial effects on the water cycle, biodiversity, and livelihood of the local population [8]. This concern is reinforced by increasing evidence that the rate of warming is magnified with elevation, with ecosystems at high altitude (> 4000 m) showing more rapid changes in temperature than those at lower elevation [4]. In addition to warming of mean temperatures, there is often a greater increase in the daily temperature variations [9] with potential effects on wildlife and humans.

Overgrazing disturbs plant species composition, biomass and vegetation characteristics [10]. These changes can lead to specified selective palatability issues for grazing animals and changes in nutrient availability [11]. At higher elevations, the climatic zone does not permit tillage crop farming. Therefore animals provide food to an increasingly dense human population in this area of Pakistan by transforming grass and herbage into milk and meat [5]. In recent years larger herds of cows, horses, and sheep have been introduced in many high-altitude regions over the world [12,13], which previously did not support large grazing herds [14,15]. Changes in the agricultural system by the introduction of new crops and the replacement of native species of grazers by species from other environments could be complex in the removal of certain endogenous plants as well as overall plant cover reduction. Warming and grazing both have substantial impacts on the natural elements of high-altitude ecosystems, with potentially vital consequences for local livelihoods [16]. In cold habitats such as the high Himalayas, plant leaves are generally rich in nutrients to counteract the negative effects on plant growth induced by this harsh environment [17] and are therefore highly palatable for cattle. For instance, as a result of extensive surveys to evaluate the nutritional status of higher altitude plants, it was proposed that higher altitude plants always had higher N content per unit leaf area when compare with contrasting altitudes a this increased with altitude in herbaceous plants [18]. Beyond grazing, one of the greatest services provided by plant communities to indigenous people of mountainous regions is their use for traditional medicine. The therapeutic effect of some mountain plant species is well known in many regions [12,19–22], yet detailed ethnobotanical studies are scarce [23].

While a great deal of research has been conducted to predict warming and overgrazing effects on alpine biodiversity, there is an important need to combine traditional indigenous knowledge with modern scientific knowledge [20]. In particular, these predictions have yet to incorporate perspectives that assess threats to the linked biological and cultural systems of local people [24–26]. Here, we provide an example of integrating plant community responses to grazing and warming with benefit-relevant indicators to assess how people’s access to culturally important plant species may be affected in the future.

This study was performed in Khunjerab National Park (KNP) located in the Pakistani Himalayas. This region is interesting to study the combined effect of warming and grazing on culturally important plant communities for several reasons. First, the medicinal plants of the Eastern Himalaya are an invaluable resource for the local communities and also Pakistan [27]. Second scientists have been gathering climatic data from this remote area for over 50 years, indicating that the average temperatures have increased by 1.5 °C, more than twice the global average increase (World Economic Forum 2020). Third, the mountainous areas of KNP are extensively used as feeding grounds for both wild ungulates (yaks) and domestic animals (goats, sheep, and cattle) with most families keep mixed herds [29]. Livestock pressure is progressing as the human population is growing as animal production is the primary means of productivity. Merging traditional knowledge with manipulative experiments is a helpful approach to test the sensitivity of such ecosystems to climate change [17]. The specific objectives of this study were to 1) document the indigenous knowledge of culturally important medicinal plants and their occurrence through ethnobotanical surveys; 2) quantify the effects on these plants of climate change (warming) and grazing through manipulative experimentations.

## Materials and Methods

### Study Area

The current study was conducted at the Khunjerab National Park (KNP) which is situated in the Hindukush-Himalaya (HKH) mountain ranges near the border with China (36.37° N, 74.41°E). The KNP is spread over an area of 4455 km^2^ with altitudes ranging from 2439m to 4878m above sea level. [30,31]. The climate is defined by warm summers starting from May and it lasts till early August at some lower altitudes (3340m) while at higher altitudes it ends in late July Winters are harsh and cold. The maximum temperature recorded in May goes up to 25°C in the warm year [32] while in winter it drops down below 0°C from October (3590m) [15,33,34]. Annual precipitation ranges from 200 to 900 mm per year [31,35]. Being alpine KNP harbors the following vegetation altitudinal zones: 1) The *sub nival zone (> 4500 m)* is made of snow and bare desert, covering about 30% of the park’s area. Characteristic plant species are *Saussurea simpsoniana, Primula macrophylla Oxytropis microphylla, Potentilla desertorum*, and *Hedinia tibetica* [36]. 2) *Alpine meadows (3500–4500 m)* are rich in herb biomass and therefore serve as important habitat for livestock (sheep, goats, cattle’s and yoks) and wild herbivores such as ibex (*Capra ibex sibrica)*, golden marmot (*Marmota caudata aurea*), and Marco Polo sheep (*Ovis ammon polii*). Dominant plant taxa are *Primulla macrophylla, Plantago lanceolate, Saxifraga* sp, *Potentilla multifida, Poa alpine* and *Carex spp*. 3) The *sub alpine Steppe (< 3500m)* is vegetated mainly with *Artemisia* and *Primula* plant genera. Some grasses such as *Poa* and *Carex* sp. are also found in relatively moist places [37]. The tremendous increase in the population of inland herbivores (> 5026) and their reliance on rangelands has placed immense burden on the highland rangelands [38].

The HKH region harbors a rich indigenous knowledge which serves as a source of sustainable rangeland management [24]. The region is also home to the most versatile cultures, languages, traditional wisdom, and religions. The significant aspect of livelihood in HKH is inherent to mountain specificities (limited accessibility, unique richness, greater fragility socioeconomic inequalities, indigenous knowledge, and vulnerability). It is home to native people who are most marginalized socially and economically and vulnerable to ongoing environmental changes [20,39,40], in particular global warming [1,41–43], [44,45]. Traditional knowledge may help local people to find solutions helping them to cope with impending changes [42,46,47].

### Ethnobotanical Surveys

We designed questionnaires to obtain information about how accessible and/or useful are medicinal plant species for local people in the HKH region. We visited three distinct valleys of KNP (Khunjerab, Ghunjerab, Shimshal) and negotiated with local community coordinators to enlist the people interested in the survey. Socio-economic and demographic data about interviewed people are given in supplementary S2 Table. Partially structured interviews were formulated to collect information about key medicinal properties of plants, frequency of use, and the perceived impacts of climate change on their life cycle, growing period, and occurrence. The composed ethnomedicinal data was quantitatively reviewed by the index of Relative Frequency Citation (RFC), calculated as the ratio between the number of informants mentioning the use of a given species and the total number of informants participating in the survey [37].

### Warming and Grazing Experiments

As part of the Multi-Site Experiment in Rangeland and Grassland Ecosystem (MERGE) project, a three-year experiment was established in 2015 and visited biannually year for maintenance and collection of data the observations. The project included the installation of an experimental set up at five different sites along an altitudinal gradient ranging from 3590 to 4696 m (Fig 1a). Within each fenced area, we selected two plots with similar slope and soil features. Each site consisted of a fenced area (un-grazed) of 5 × 5 m, which was designed as a randomized block design (RBD) and laid out four subplots of 2.5 × 2.5 m. Among those, we selected two subplots for warming treatment using fiberglass open-top chambers (OTC) (Fig 1b). They were hexagonal with a 32 cm height having an outer edge of 1.5 × 1.5 m and a central portion of 1 × 1 m open at the top [48]. The OTCs were kept on the subplots in each summer season. The warming effect of OTC at 10 cm above the soil surface was between 1.7–2.3°C on average. All recordings of plant community composition and abundance were performed during the peak blooming season (Fig 1d) which starts from late March and lasts until late August [15]. Two subplots (grazing with manual clipping and control (no warming) were considered to employ grazing treatment. Inside the fence, grazing treatment was simulated by manual clipping of the vegetation to the average height of plants eaten by herbivores outside. Clipping was done during the peak season (June–Aug). The grazing levels were permanently marked at the starting year and each plant species growth was recorded individually from the clipped point.

**Fig 1.**
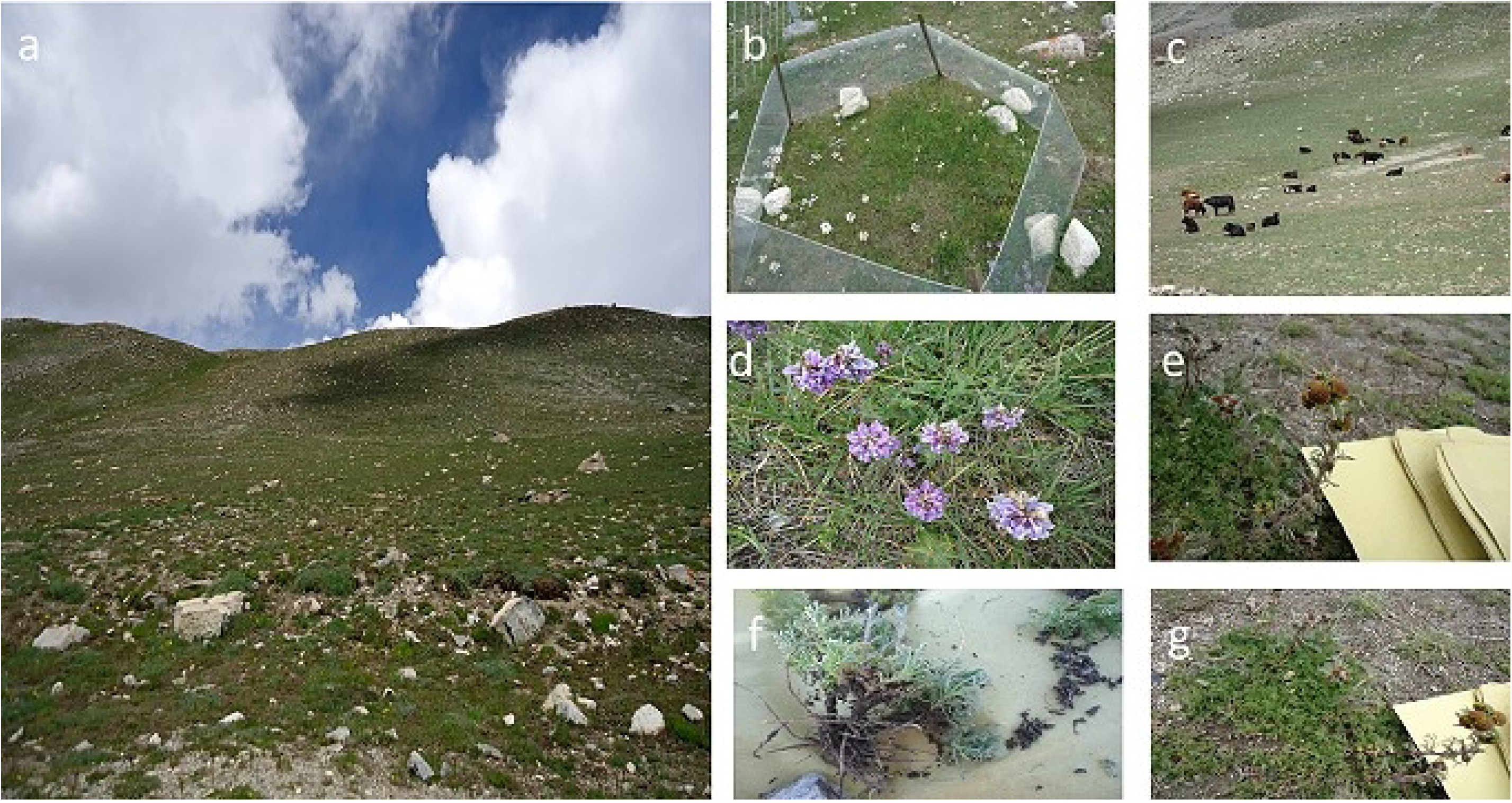
Landscape of the study region and the Multisite Experimentation at Rangeland and Grassland Ecosystem (MERGE) site at Khunjerab National Park (KNP), Pakistan. **(a)** pastures used for grazing herds of local communities. **(b)** Warming experimental setup with *Silene gonosperma* inside an open top chamber (OTC). (c) Some livestock herds (yolks, sheeps) grazing on the pastures near the experimental site in July 2017. **(d)** *Astragulus penduncularis* in full bloom at the grazing treatment. **(e)** Inventorying field observations of *Artemisia rupestris*. **(f-g)** Plant collection.

**Fig 2.**
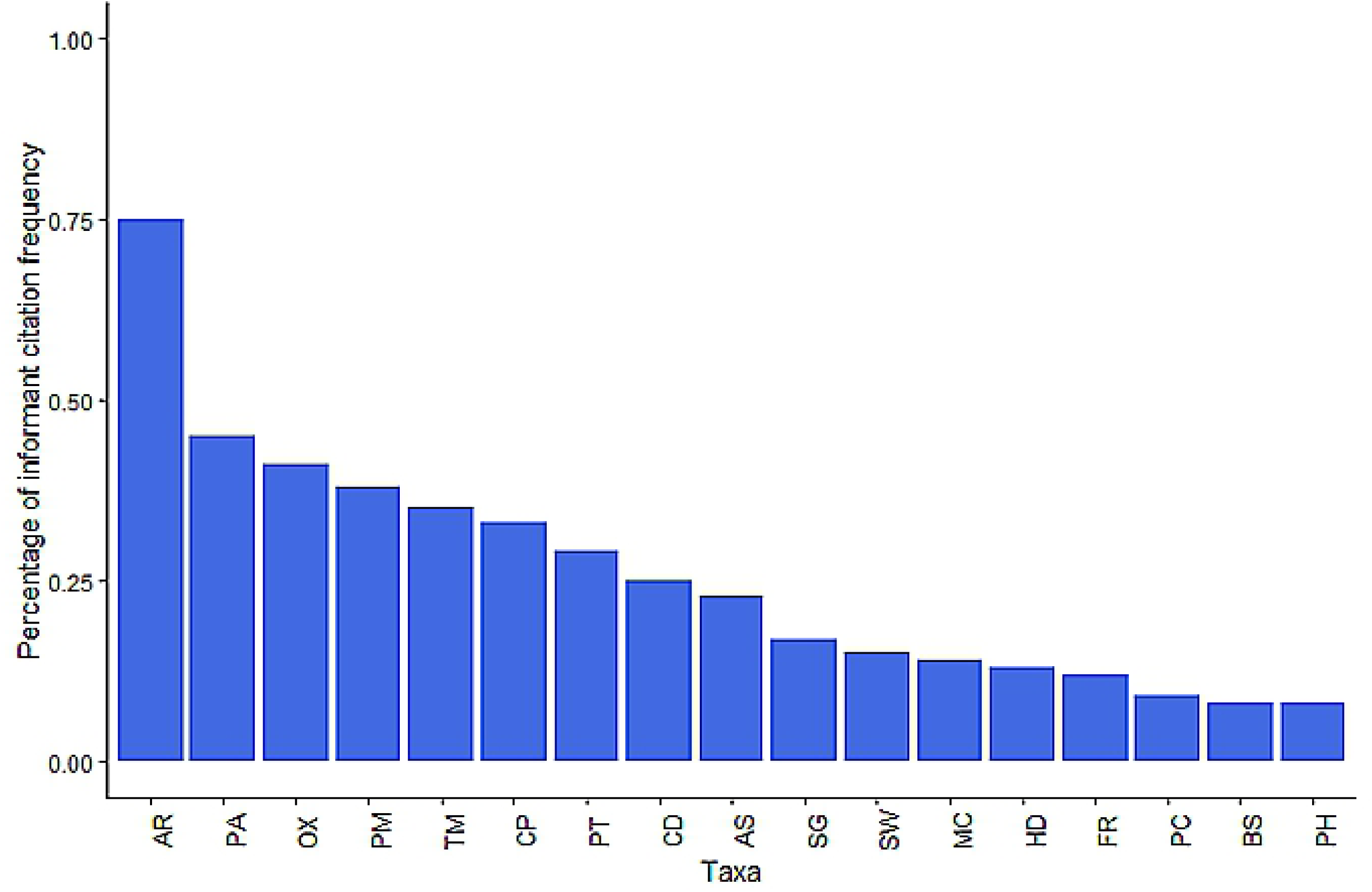
Percentage of Informant Citation Frequency Concerning the Medicinally and Culturally Important Species in KNP. The frequency is expressed as percentage a plant is mentioned by the informants. \**Artemisia rupestris* (AR), *Poa alpine* (PA), *Oxytropis glabra* (OX), *Plantago major* (PM), *Tamaricaria officinalis* (TM), *Comastoma pulmonarium* (CP), *Potentilla hololeucaI* (PT), *Carex divisa* (CP), *Astragulus penduncularis* (AS), *Silene gonospermum* (SG), *Smelowskia calycina* (SG), *Primula macrophylla* (MC), *Hedinia tibetica* (HD), *Saxifraga* (SG), *Pedicularis kashmiriana* (PC), *Bistorta officinalis* (BT), *Peganum hermala* (PH).

### Data Sampling and Analysis

During the experimental years (2016–2018), we used the quadrat sampling method [49] to measure plant metrics such as percentage cover, plant height, and aboveground biomass. The percentage cover of the selected plant species was recorded individually at the start of the experiment (2016) by visualizing the numbers of grids of quadrat covered by a plant species. In each 1 × 1 m subplot, the percent cover for each plant species was estimated to the nearest 5% for each species rooted inside the plot using the Daubenmire method [50]. Plant height of at least five individuals of each species was measured in both the OTCs and control plots, as the distance between the ground and the highest photosynthetic part. In all experimental sites, individual plant species were identified and sampled carefully for taxonomic classification (Fig 1f). Inventory was updated each year to record the new observations (2016–2018). We clipped the vegetation from OTC and the above-ground biomass of the plant species was measured by air drying samples at 37°C for 72 hours.

We conducted one–way ANOVA to examine the difference between percentage cover and biomass of plant species under warming and grazing treatments. A Post-hoc Tukey test was applied to compare the responses of plant species to a specific treatment (S3 Table a&b). To assess the change in the relative distribution of plant species within the plant communities we used non-metric multidimensional scaling (NMDS) ordination analyses [51] on plant species that were present across the plot at each site (3590m–4696m) so that we could characterize plant community changes for each warming and grazing treatment groups (see supplementary S2 Fig.). We used Bray–Curtis dissimilarity measure to deal with relative abundance data as it allows using both presence/absence and abundance data. The vegan package R (metaMDS) was used for analysis. All data analyses were carried out using R version 3.5.3 (2019).

This study is part of Ph.D. research project titled “Developing strategies and baseline protocols to predict climate change impacts on selected plants and prevelance of associated allergic diseases in Pakistan”. Here it is confirmed that this specific study involved plant material and interviewed people of the study area. The cosent of agreement of the interviewed people was take verbally by 1^st^ and 2^nd^ author. The work has been performed at all times with ethical oversight by an ethics committee and was reviewed and approved prior to start of the work, by Institutional Ethics Review Board (No. CIIT/BIO/ERB/17/53) and Institutional Biosafety Committee (No. CIIT/BIO/ERB/17/04) of COMSATS Institute of Information Technology (CIIT) Islamabad.

## Results

### Ethnobotanical Survey

We interviewed a total of 80 informants who identified approximately 50 medicinal plant species widely spread in the area (see supplementary S1 Table). Those interviews also allowed us to collect detailed information about the therapeutic properties of plant species along with their altitudinal range of occurrence, time of blooming, part used for remedies, and potential climate change effect on their availability (Table 1). Overall, people use plants to treat various diseases such as high blood pressure, stomachache, wounds, cold/fever, rheumatism, asthma, diarrhea, hepatitis, and diabetes (see supplementary S1 Fig.).

**Table 1.** Ecological and Ethnobotanical Information on the Plant Species Assessed During the Study Period.

| Plant species and authority | Family | Local name | Altitudinal range (m) | Period of availability | Part used | Medicinal properties | Climate change effect (according to local people) | RFC* |
| --- | --- | --- | --- | --- | --- | --- | --- | --- |
| *Artemisia santolinifolia* Turcz.ex Krasch | Asteraceae | Roon | 3343-4039 | Whole year | Whole plant | Worm, digestion, diarrhea, malaria | Increased availability due to extended growth period | 0.9375 |
| * *Artemisia rupestris* L. | Asteraceae | Khich | 4059-4676 | Whole year | Whole plant | Digestion, insect repellent elevates immunity, skin diseases | Late blooming | 0.9375 |
| * *Taraxacum officinale* L. | Asteraceae | Yamook | 3346-4059 | Whole year | Stems, leaves, dried twigs | Asthma, cardiac diseases | No effect | 0.4375 |
| *Smelowskia calycina* (Steph.) | Brassicaceae | Zakh | 4096 | July- Oct | Flower, leaves | Blood pressure troubles and diabetes | Berries are not that juicy | 0.1875 |
| *Silene gonosperma* (Rupr.)Bocquet | Caryophyllaceae | Gulch | 3359-4693 | May-Aug | Whole plant | Fever | Vegetation has increased | 0.2125 |
| * *Astragalus penduncularis* Royle ex Benth | Fabaceae | Zhoop | 3346-4696 | May-July | Leaves and Flower | Diabetes, antiaging, anti-inflammatory | Availability increased | 0.2875 |
| *Oxytropis glabra* DC | Fabaceae | Zarth sprag | 4034 | March-Aug | Leaves | Internal wound, anti-aging | No effect | 0.5125 |
| * *Comastoma pulmonarium* (Turcz) Toyok | Gentianaceae | Shalay char | 4696 | May-Au | Leaves and flower | Pneumonia, sore throat and fever. | Not observed | 0.4125 |
| *Peganum harmala* L. | Nitrariaceae | Ispandur | 3343 | May-Jul | Seeds | Fever and joint pain | No effect | 0.1 |
| *Plantago major* L | Plantaginaceae | Sepgilk | 4059 | May-Sep | Leaves and Seeds | Ulcers, skin wounds & dysentery | Ripe late | 0.475 |
| * *Poa alpina* L. | Poaceae | Noz | 3346-4076 | Mar-Oct | Whole plant |  | Denser vegetation | 0.5625 |
| *Bistorta officinalis* | Polygonaceae | onbu | 4690 | May-Aug | Root | Wounds | Less vegetation | 0.1 |
| * *Primula macrophylla* D.Don | Primulaceae | Banafsha | 4676 | Whole year | Leaves and Flower | Alleviate pain, analgesic | Forage available | 0.175 |
| * *Potentilla hololeuca* (Boiss)Hook.f | Rosacea | Zatsprig | 3343-4696 | May-Jul | Leaves and Flower | Menstrual cramps | No effect | 0.3625 |
| *Saxifraga* sp | Saxifragaceae | Sitbark | 4693 | June-Aug | Leaves | Diarrhea, and minor skin problems | No effect | 0.15 |
| *Pedicularis kashmiriana* Pennell | Scrophulariaceae | Push | 4690 | Jun-Aug | Whole plant | Stomach aches | Shifted towards high altitude | 0.1125 |
| *Hedinia tibetica* (Thomson) Oestenf | Scrophulariaceae | tibet | 4690 | Jun-Aug | Whole plant | Pain and swelling, muscle weakness | Range shift from low altitude towards high | 0.1625 |

(\*) represents most preferable forage species, RFC: Relative frequency citation indicates *Artemisia species* having a relatively high frequency of citation among other species

### Culturally Important Plant Response to Warming and Grazing

Experimental warming had an overall positive effect on the biomass of most plant species (+1.3% on average), yet this trend was not significant due to the large inter-site variability in species responses (Fig. 3a, see supplementary S3 Table a). Instead, vegetation cover increased significantly in warmed plots, on average by 5.5% as compared to control plots (Fig 3b, S3 Table b). Plant cover increase in response to warming was highly variable among taxa, ranging from 1 ± 0.6% for *Bistorta officinalis* to 18.7 ± 4.2% *for P. alpina* (Fig 3b). Overall, our NMDS analysis revealed that plant communities showed higher differences among sites than among warming treatments, reinforcing the idea that plant response was dependent on altitude (see supplementary S2 Fig.).

**Fig 3.**
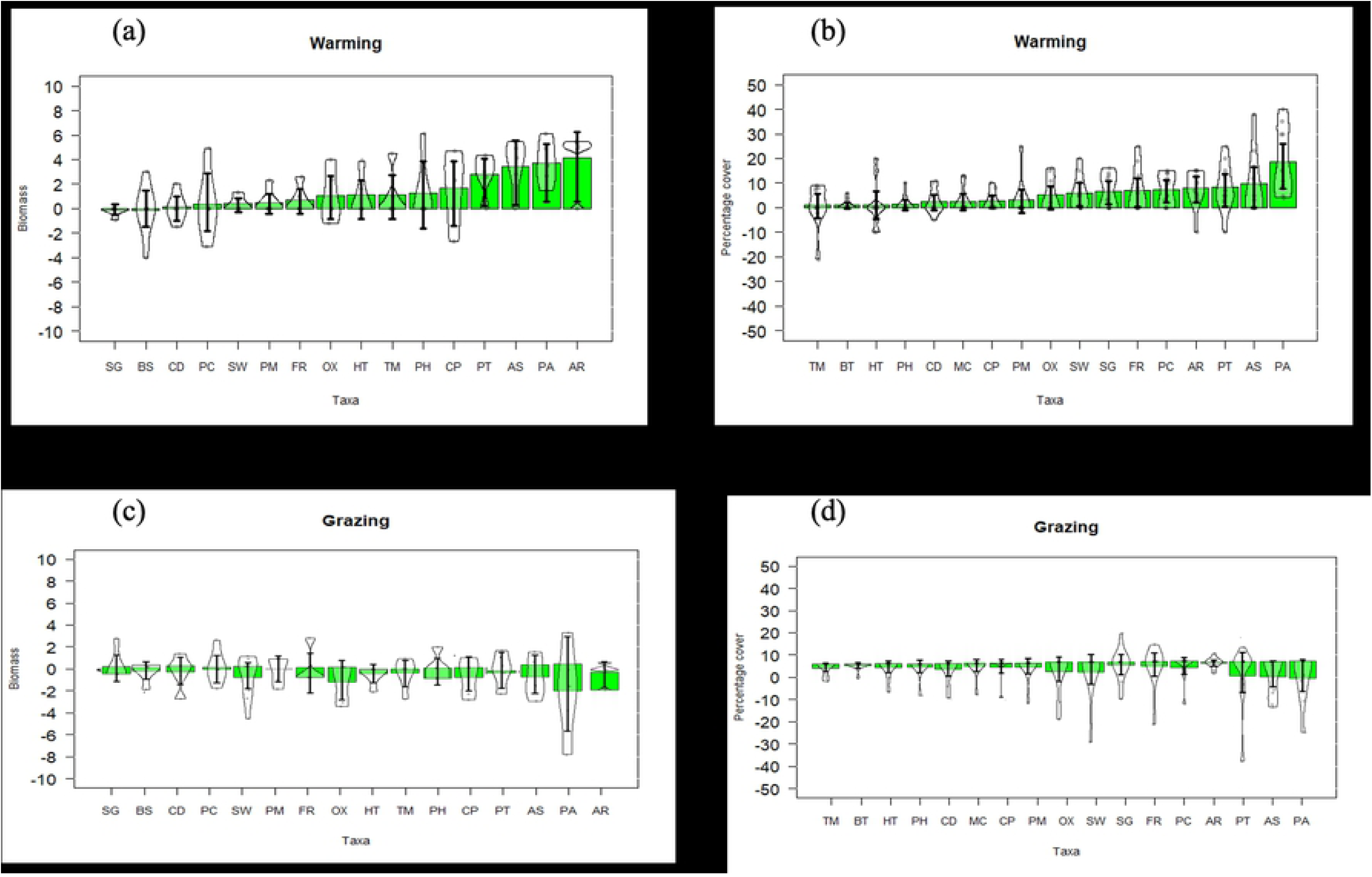
Effects of warming and grazing treatments on the biomass and percentage cover of plant species during the experimental year 2016–2018. The data represents mean±standard errors of all plant species. **(a)** shows the warming effect of biomass. **(b)** shows the warming effect on percentage cover. **(c)** shows the grazing effect on biomass. **(d)** shows the grazing effect on percentage cover. Different letters denote statistically significant differences as calculated by post hoc Tukey’s test at P < 0.05, *Artemisia rupestris* (AR), *Poa alpine* (PA), *Oxytropis glabra* (OX), *Plantago major* (PM), *Tamaricaria officinalis* (TM), *Comastoma pulmonarium* (CP), *Potentilla hololeucaI* (PT), *Carex divisa* (CD), *Astragulus penduncularis* (AS), *Silene gonospermum* (SG), *Smelowskia calycina* (SW), *Primula macrophylla* (MC), *Hedinia tibetica* (HD), *Saxifraga* (FR), *Pedicularis kashmiriana* (PC), *Bistorta officinalis* (BS), *Peganum hermala* (PH).

Compared to the control plots, grazing treatments had an overall negative effect on plant biomass and percentage cover (Fig 3c & 3d). Yet, once again, this general trend was not significant due to the high variability in plant response among sites. The extent of increase ranged from 0.1 ± 0.5% in *Carex divisa* to 4.14 ± 1.04 % in *A. rupestris*.

We further assessed the response to experimental warming and grazing for the five plant species that were most cited by surveyed people (*A. rupestris, Astragulus penduncularis, P*.*alpina*, P. hololeuca and P macrophylla). These responses were divided into four categories: combined positive effects of warming and grazing (+W, +G), combined negative effects of warming and grazing (–W, –G) and antagonistic effects (–W, +G or +W, –G) (Fig 4, Table 1). The mean percentage cover of species was affected positively by warming and negatively by grazing (Fig 4, +W, –G zone). One species, A. rupestris, was favored by both warming and grazing. However, for P. alpina and P. hololeuca, increased in cover as a response to warming at some sites made them more susceptible to grazing. P. macrophylla did not show any significant response to either grazing or warming except at one site (aggressively grazed) where it was negatively affected by both treatments (Fig 4).

**Fig 4.**
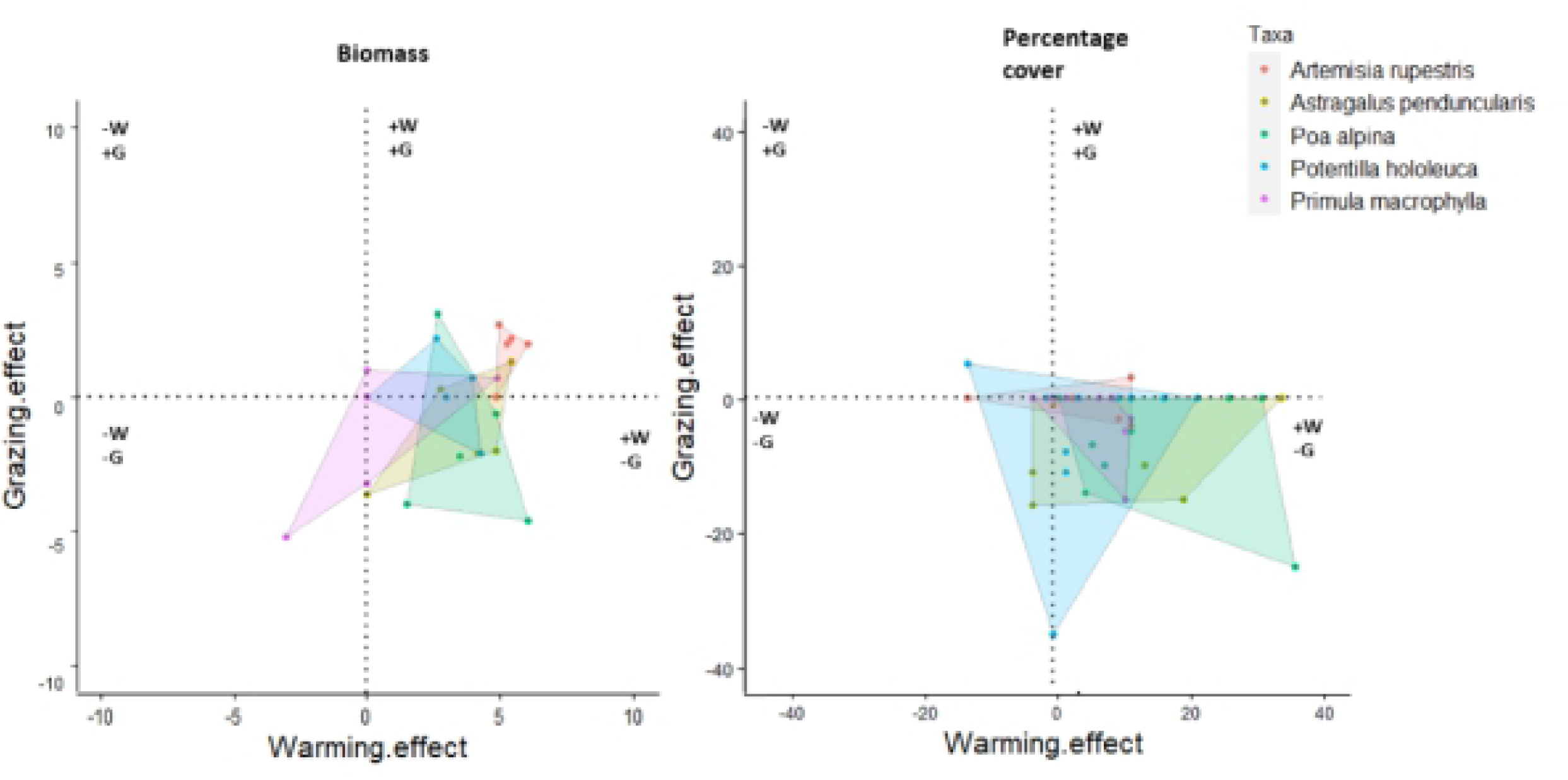
Susceptibility Zones of Five Most Frequent and Culturally Important Plant Species into Under Combined Impacts of Warming (W) and Grazing (G) Treatments. The susceptibility zones are combined with positive effects (+W, +G), combined negative effects (–W, – G), and antagonistic effects (–W, +G or +W, –G). Polygons join the dots of the presence of a species in the respective zone. **(a)** Biomass **(b)** Percentage cover.

## Discussion

According to the Intergovernmental Panel for Climate Change [52] indigenous, local and traditional knowledge systems and practices, including indigenous people holistic view of community and environment, are a major resource for adapting to climate change. HKH culture and livelihoods depend on local medicinal plants for medicine, food, grazing, wood, etc. and several studies have shown the importance of traditional plant use for indigenous people in the Himalayas [15,33,37,53]. Our study confirms that the HKH region is rich in indigenous knowledge about culturally important plants that are used for various human diseases (Table 1, Fig. 2) as well as livestock fodder and forage. For example, local mountain communities tend to graze their animals on selective nutritious plant species [54]. Our survey revealed that local people had extensive knowledge about plant identity, occurrence in the area, population trends in recent years, and their medicinal and nutritional properties. Plant species from *Asteraceae* (e.g. *Artemisia rupestris, Artemisia santinofolia, Artemisa rutifolia*) were particularly important for the family medication (cited by 25% of respondents) as reported in previous studies [30,33]. For example, a whole plant decoction of *Artemisia* is used for the treatment of fever, stomach pain, vomiting, and skin ulcers, under both pharmacological and clinical information [55–57]. Interestingly, our study revealed that the *Asteraceae* family had the highest RFC value (0.9) due to their increased availability in warming treatments. This plant group may, therefore, be an ally for people facing global warming. Interestingly, our study revealed that the *Asteraceae* family had the highest RFC value (0.9) due to their increased availability in warming treatments. This plant group may, therefore, be an ally for people facing global warming.

The impact of local climate change on traditional livelihood is strong and evident but can be analyzed for both positive and negative aspects. Warming may have positive impacts on overall vegetation richness and productivity in cold areas [2,58–60] as supported by our results. Previous warming experiments and studies carried out to assess the effect of temperature on alpine ecosystems using OTCs generally concluded that warming enhances the plant vegetation cover and height [21,39,48,61]. This pattern is consistent with our results that showed a positive effect of warming on plant community biomass. Such an increase in biomass was highly taxa-specific, favoring some species upon others, and thereby modifying community composition. This taxa-specific pattern of plant community responses to warming has been stressed in other studies performed in challenging environmental conditions such as the Arctic [11,35,61–63]. Flower production response to warming was positive in 41% of plants but had no effects on the other taxa. Interestingly, our result further showed that *P. alpina, A. rupestris, A. penducularis, P. multifida, and P. macrophylla* can survive in warmer conditions at high elevations, suggesting that they may expand their range if herbivory is controlled. This reinforces the idea that in some cases, warming may generate new opportunities for people living under cold conditions [64]. While previous studies suggested that overgrazing is one of the major drivers of rangeland degradation [18,65– 70]. Our results only partly supported this assumption, while grazing treatments did have an overall negative effect on plant biomass and percentage cover, the general trend was not significant due to the high variability in plant response among the sites. This may be explained by the high heterogeneity in cattle spatial distribution at the landscape scale. Except for two highly palatable species (*P. alpina* and *A. rupestris*, the percent cover of most culturally important plants in KNP were not significantly lower outside cattle exclusion plots, suggesting that coordinated management of cattle herds among local communities may help in keeping the negative impact of grazing at sustainable levels. A constant increase in species covers though the time can occur under moderate grazing conditions [65,71] thus assuring the availability of culturally and medicinally important plant species.

## Conclusions

Overall, our study is the first to provide experimental evidence at Khunjerab National Park, of the combined effect of warming and grazing on culturally important plants. Our results provide valuable information for the evaluation and prediction of grasslands sensitivity to future threats. Local knowledge can be rapidly and efficiently gathered using systematic tools, and it can provide a jumping pad to policymakers for designing mitigation and adaptation strategies for climate change in a region that is undergoing rapid change and for which scientific data are meager. However, the prevailing indigenous knowledge in the study area is facing an uncertain future. As an example, the nature of traditional knowledge is making it more difficult to learn and then transfer it accurately. Furthermore, practicing traditional therapies are not being respected by new generations. Other challenges include low literacy rate in the study area, no proper documentation of indigenous knowledge, and the introduction of modern allopathic medicines, rapid technological advancement, and environmental degradation.

## Acknowledgments

We are highly thankful to local influential persons, hunters, and guides who facilitated and supported the survey team. The English language of the manuscript has been improved by Prof. Dr. Richard Goodman which is acknowledged.

## Authors Contributions

SK conducted the practical work, interviewed the people, compiled the literature, and wrote the manuscript. MN helped SK in the practical work execution, interviewed the people and data collection from KNP. SN helped in manuscript organization, and improvmed of the final manuscript. AA reviewed the manuscript. OD helped in data compilation & statistical analyses and manuscript writing. ZA designed the strategy, supervised the overall research work, and improved the manuscript.

## Supporting Information

**S1 Table docx. Plant Species Composition at MERGE Experimental Sites**

A variety of plant species in experimental sites at different elevations. Each site was catagorised according to the number of species present.majority of selected species were present at each site, but there was a representative specie of each elevation. *Bistorta officnalis* is present only at highest altitude(4696m), similarly *Plantago major* (3690m) is present on site 5, lower altitude

**S2 Table docx. Specific Characteristic of Local Informants**

Demographic features of informants interviewed for ethnobotanical information of culturally important plants. In the survey, their gender, age and socioeconomical details were recorded.

**S3 Table A. docx. Warming Effect on Aboveground Biomass of Plant Species**

Data represents the mean±SE of various plant species biomass in response to warming effect. Means followed by similar letter are not significantly different form each other as determined by Post-Hoc Tukey’s at p<0.05.warming overall increased the above ground biomass of plant species but this response was taxa se

**S3 Table b docx. Warming Effect on Percentage Cover of Plant Species**

Data represents the mean±SE of various plant species percentage cover in response to warming effect. Means followed by different letters are significantly different form each other as determined by Post-Hoc Tukey’s at p<0.05.

**S1 Fig pdf. Ethnobotanical dominant families and treated ailments**

(A) percentage of ailments treated by understudy medicinal plant families. (b) percentage of dominant culturally important plant families present in the study area

**S2 Fig pdf. Changes in the distribution and occurrence of plant species in response to climate warming over a 3–years experimental period along the elevation gradient of (3590–4696m)**.

The relative abundance of plant species (presence/absence) shown by paths of mean values in Non-metric multi-dimensional scaling (NMDS) using Bray and Curtis dissimilarity index in R. OTC = Open top chamber, OTC1, OTC2, two OTC per site

Site 1 4,696m near china boarder, Site 2= 4,059m Site 3= 4022m Site4=3,990m Site 5= 3,590m

